# Bacterial tail anchors can target to the mitochondrial outer membrane

**DOI:** 10.1101/111716

**Authors:** Güleycan Lutfullahoğlu Bal, Abdurrahman Keskin, Ayşe Bengisu Seferoğlu, Cory D. Dunn

## Abstract

During the generation and evolution of the eukaryotic cell, a proteobacterial endosymbiont was refashioned into the mitochondrion, an organelle that appears to have been present in the ancestor of all present-day eukaryotes. Mitochondria harbor proteomes derived from coding information located both inside and outside the organelle, and the rate-limiting step toward the formation of eukaryotic cells may have been development of an import apparatus allowing protein entry to mitochondria. Currently, a widely conserved translocon allows proteins to pass from the cytosol into mitochondria, but how proteins encoded outside of mitochondria were first directed to these organelles at the dawn of eukaryogenesis is not clear. Because several proteins targeted by a carboxyl-terminal tail anchor (TA) appear to have the ability to insert spontaneously into the mitochondrial outer membrane (OM), it is possible that self-inserting, tail-anchored polypeptides obtained from bacteria might have formed the first gate allowing proteins to access mitochondria from the cytosol. Here, we tested whether bacterial TAs are capable of targeting to mitochondria. In a survey of proteins encoded by the proteobacterium *Escherichia coli*, predicted TA sequences were directed to specific subcellular locations within the yeast *Saccharomyces cerevisiae*. Importantly, TAs obtained from DUF883 family members ElaB and YqjD were abundantly localized to and inserted at the mitochondrial OM. Our results support the notion that eukaryotic cells are able to utilize membrane-targeting signals present in bacterial proteins obtained by lateral gene transfer, and our findings make plausible a model in which mitochondrial protein translocation was first driven by tail-anchored proteins.

## BACKGROUND

During the integration of an α-proteobacterial endosymbiont within the eukaryotic cell, genes transferred to the (proto)nucleus were re-targeted to mitochondria, allowing these organelles to remain the location of crucial cellular processes [1-3]. In addition, other polypeptides that evolved within the eukaryotic lineage or that were acquired through lateral gene transfer from other organisms were directed to mitochondria [4-6]. Across eukaryotes, the β-barrel Tom40 protein forms a pore by which proteins pass through the OM [7-9]. However, the Tom40 polypeptide seems to require already existing TOM complexes for mitochondrial insertion [10,11], giving rise to a “chicken or the egg” dilemma when considering how the TOM complex may have evolved.

Several narratives might be proposed for how mitochondria first evolved the ability to transport proteins from the cytosol. In one scenario, an early translocation pore that was self-inserting at the mitochondrial surface might have allowed mitochondria to begin to import proteins, permitting the subsequent evolution of the translocon found in eukaryotes today [12]. Current evidence suggests that the self-insertion of tail-anchored proteins at the mitochondrial OM is possible [13-15], and some tail-anchored pro-apoptotic proteins appear to have the ability to generate membrane pores at mitochondria [16,17], making tenable such a scenario for the evolution of mitochondrial protein import. At the inception of mitochondria, such tail-anchored proteins would likely have been derived from prokaryotes, particularly if mitochondria were required for the generation of the stereotypical compartmentalized structure of eukaryotes.

We focused our attention upon a single aspect of this hypothesis: can TAs obtained from bacterial proteins be inserted into the mitochondrial OM when expressed within a eukaryotic cell? Indeed, our results demonstrate insertion and function at the mitochondrial OM for predicted TAs encoded by the proteobacterium *E. coli*, and we describe the relevance of our findings to the concept of lateral gene transfer during eukaryogenesis.

## RESULTS

### Bacterial Tail Anchors Can Localize to Mitochondria

To test whether predicted bacterial TAs might have the capacity to be inserted at the mitochondrial OM, we identified 12 *E. coli* proteins predicted to harbor a solitary α-helical transmembrane (TM) domain at the polypeptide carboxyl-terminus (Fig. S1), then fused mCherry to the amino-terminus of these TAs and examined their location in *S. cerevisiae* cells by fluorescence microscopy. mCherry-ElaB(TA) (Fig. 1A) and mCherry-YqjD(TA) (Fig. 1B) were readily detectable at mitochondria, as reported by co-localization with superfolder GFP (sfGFP) [18] fused to the TA of the *S. cerevisiae* Fis1 polypeptide, a protein playing a role in yeast mitochondrial division. A lesser fraction of mCherry-ElaB(TA) and mCherry-YqjD(TA) was localized to the endoplasmic reticulum (Fig. S2). ElaB and YqjD are members of the DUF883 family of proteins. Little is known about the function of DUF883 family members, but YqjD may recruit ribosomes to the *E. coli* plasma membrane during stationary phase [19]. Although negligible fluorescent signal was detectable by microscopy or flow cytometry (C. Dunn, unpublished results), mCherry-TcdA(TA) could also be visualized at mitochondria (Fig. S3A). TcdA (also called CsdL) catalyzes the modification of *E. coli* tRNAs [20].

**Figure 1.**
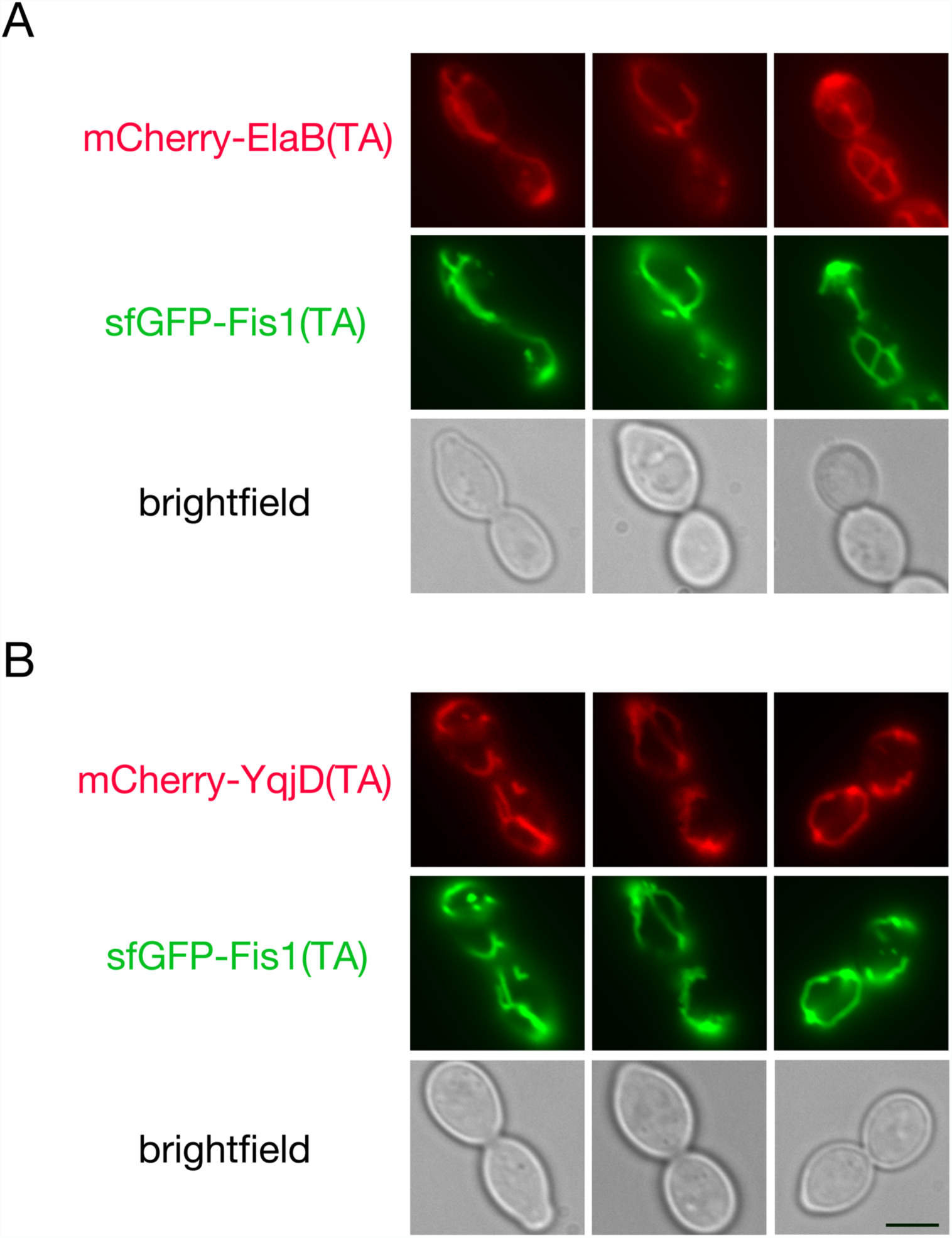
The predicted ElaB and YqjD TAs localize to mitochondria. Strain BY4741, harboring plasmid b294 (sfGFP-Fis1p), was mated to strain BY4742 carrying mCherry-ElaB(TA)-expressing plasmid b275 (A) or strain BY4742 carrying mCherry-YqjD(TA)-expressing plasmid b279 (B). The resulting diploids were visualized by fluorescence microscopy. Scale bar, 5μm.

Other predicted TAs derived from the *E. coli* Flk, YgiM, RfaJ, DjlB, FdnH, NrfR, and YmiA proteins appeared to allow at least partial localization of mCherry to various locations associated with the endomembrane system (Fig. S4). However, no convincing localization to mitochondria was apparent after fusing any of these TAs to mCherry. Moreover, mCherry-YhdV(TA) appeared to be distributed throughout cytosol and nucleus, indicating failure to target efficiently to any membrane. mCherry-YgaM(TA) was not detectable, suggesting its degradation.

### Bacterial Tail Anchors Can Insert into Membranes in a Eukaryotic Cell

Previously, we developed an assay in which membrane insertion of proteins might be examined by a proliferation-based assay [21]. In brief, the Gal4 transcription factor is linked to a protein of interest that is thought to be membrane inserted outside of the nucleus. Failure of this fusion protein to insert at its target membrane can allow the Gal4-linked fusion protein to access the nucleus and activate Gal4-responsive promoters that drive proliferation under selective conditions. As previously demonstrated [21], while a membrane-sequestered Gal4-sfGFP-Fis1 fusion protein did not lead to a proliferation defect on non-selective medium (SC-Trp), cells carrying this construct could not survive on medium requiring activation of a Gal4p-driven *HIS3* gene (SMM-His+20mM 3-AT) (Fig. 2). Deletion of the Fis1p TA, or the presence of a A144D charge substitution within the Fis1p TA, led to a failure of membrane insertion at mitochondria, translocation to the nucleus, and Gal4-dependent proliferation on selective medium. When the TA of Fis1p, a domain whose sole purpose is to allow this protein’s insertion at the mitochondrial OM [21,22], was replaced with the TA of either ElaB or YqjD, cells were unable to proliferate on medium selective for histidine synthesis, consistent with ElaB and YqjD TA insertion at the mitochondrial OM.

**Figure 2.**
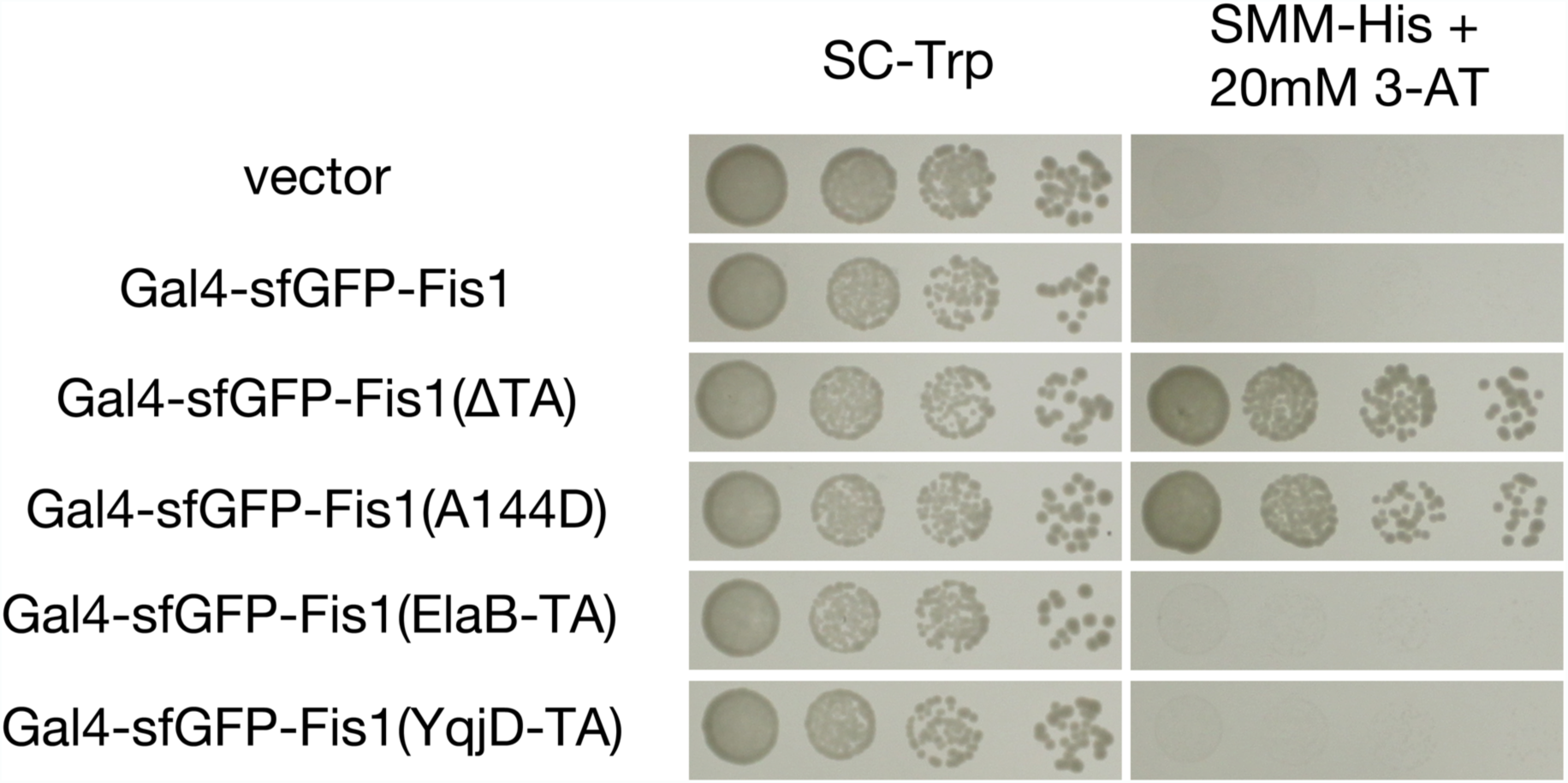
A proliferation-based assay suggests that the ElaB and YqjD TAs are membrane inserted. Strain MaV203, containing a Gal4-activated *HIS3* gene, was transformed with plasmids expressing Gal4-sfGFP-Fis1p (plasmid b100), a variant lacking the Fis1p TA (plasmid b101), a mutant containing the A144D charge substitution in its TA (b180), or the Gal4-sfGFP-Fis1p construct with the Fis1p TA replaced with that of either ElaB (b313) or YqjD (b314). MaV203 was also transformed with empty vector pKS1. Transformants were cultured in SC-Trp medium, then, following serial dilution, spotted to SC-Trp or SMM-His + 20 mM 3-AT and incubated for 2 d.

### Bacterial Tail Anchors Can Function at the Mitochondrial Outer Membrane

As these findings indicated that the ElaB and YqjD TAs may be competent for mitochondrial insertion, we tested whether these TAs can functionally replace the membrane-bound TA of Fis1p, thereby allowing Fis1p to promote mitochondrial division. Because Fis1p is required for mitochondrial fission in *S. cerevisiae*, mutants lacking this protein manifest a highly interconnected network of mitochondria due to unbalanced mitochondrial fusion [23-25]. As expected, expression of wild-type Fis1p restored normal mitochondrial distribution in this genetic background, while Fis1p prevented from mitochondrial insertion by a A144D substitution within the Fis1p TA [21] could not restore normal mitochondrial morphology (Figs. 3A and 3B). Strikingly, replacement of the Fis1p TA with the ElaB or the YqjD TA within the context of full length Fis1p polypeptide could successfully promote mitochondrial division and restore normal mitochondrial morphology. A control TA obtained from the *E. coli* YgiM protein, which is not trafficked to mitochondria, could not support Fis1p activity. In addition, a Fis1-TcdA(TA) protein could not functionally replace the Fis1p TA in this microcopy-based assay (Fig. S3B), suggesting insufficient expression, poor mitochondrial insertion, or meager functionality.

**Figure 3.**
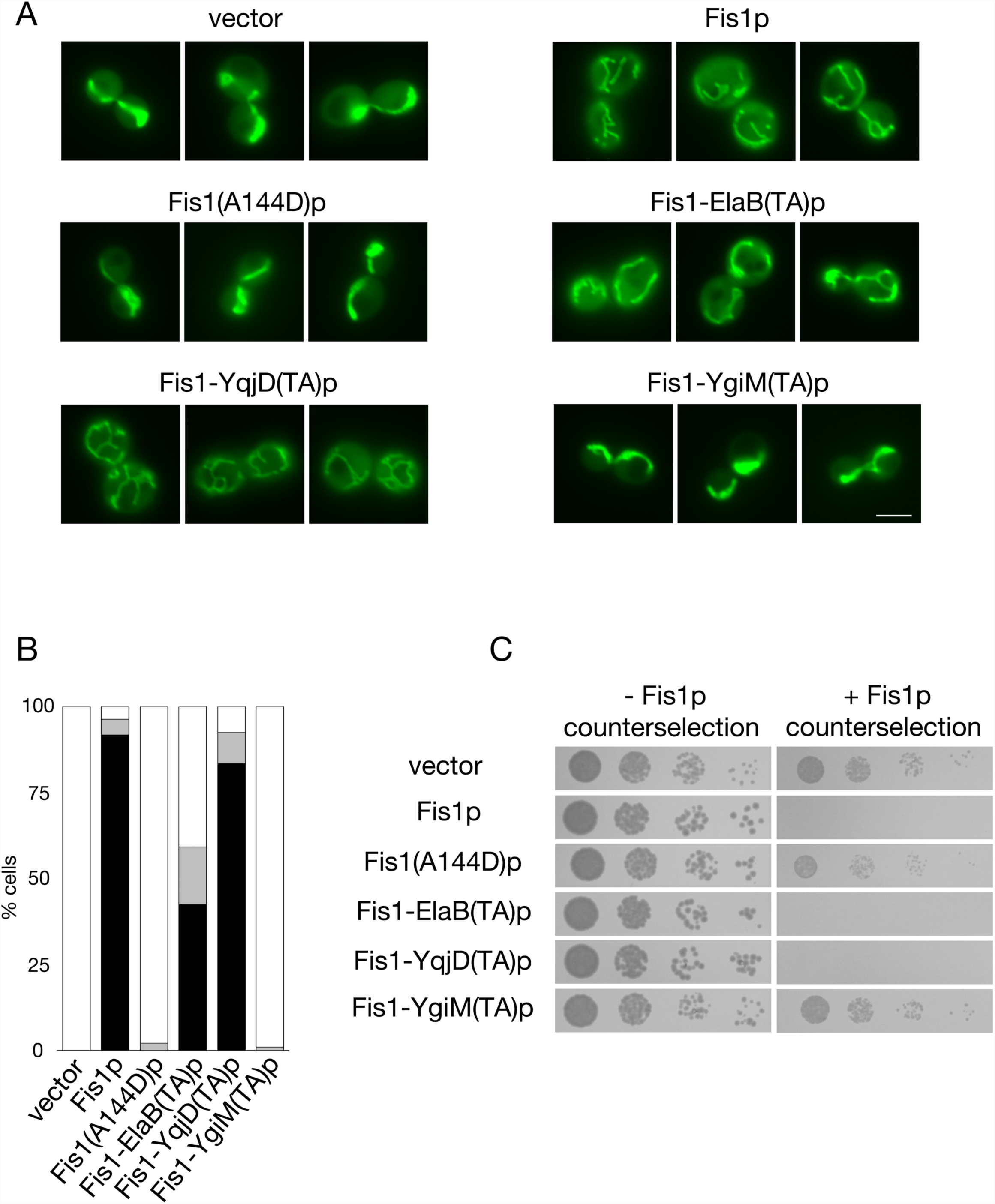
Mitochondria-localized bacterial TAs can functionally replace the TA of Fis1p. (A) The ElaB and YqjD TAs can replace the Fis1p TA in promotion of normal mitochondrial morphology. *fis1Δ* strain CDD741, expressing mitochondria-targeted GFP from plasmid pHS12, was transformed with empty vector pRS313 or plasmids expressing wild-type Fis1p (b239), Fis1(A144D)p (b244), or Fis1p with its own TA replaced by that of ElaB (b317), YqjD (b318), or YgiM (b316). Cells were examined by fluorescence microscopy. Scale bar, 5μm. (B) Quantification of mitochondrial morphology of the transformants from (A) was performed blind to genotype. White bar represents cells with fully networked mitochondria, grey bar represents cells with mitochondria not fully networked, but networked to a greater extent than wild-type cells, and black bar represents cells with normal mitochondrial morphology. Quantification was repeated three times (n>200 per genotype), and a representative experiment is shown. (C) Genetic assessment of Fis1p variant functionality. Strain CDD688 was transformed with the plasmids in (A) and proliferation was assessed without selection against Fis1p activity (YPALac medium for 2 d) and following counter-selection for cells carrying functional Fis1p (SLac-His+CHX medium for 4 d).

We then sought further evidence for functional insertion of the ElaB and YqjD TAs at the mitochondrial OM using an assay based on cell proliferation [21]. Expression of functional Fis1p in a genetic background initially lacking Fis1p and removed of the mitochondrial fusogen Fzo1p can lead to unchecked mitochondrial fragmentation, loss of functional mitochondrial DNA (mtDNA), and a corresponding abrogation of respiratory competence [26-29]. As previously reported [21], expression of wild-type Fis1p in a *fzo1Δ fis1Δ* genetic background led to an inability to proliferate on nonfermentable medium, while expression of the poorly inserted Fis1(A144D) variant did not prompt mtDNA loss (Fig. 3C). The ElaB and YqjD TAs fused to the cytosolic domain of Fis1p allowed sufficient fission activity to prompt mitochondrial genome loss from the same genetic background, again indicating successful ElaB TA and YqjD TA insertion at the mitochondrial OM. Even the Fis1-TcdA(TA) protein provoked mtDNA loss in *fzo1Δ fis1Δ* cells (Fig. S3C), suggesting some minimal level of OM insertion, and the YgiM TA again appeared unable to recruit Fis1p to mitochondria (Fig. 3C). Together, our results demonstrate insertion of the bacterial ElaB and YqjD TAs at the mitochondrial surface of a eukaryotic cell.

## DISCUSSION

Our findings, in which several predicted TAs obtained from *E. coli* can target to and function at the mitochondrial OM of *S. cerevisiae*, make plausible a scenario in which tail-anchored bacterial proteins contributed to the formation of the earliest mitochondrial translocon. The structural characteristics of the TAs of ElaB and YqjD, a helical TM domain rich in glycines followed by a positively charged patch ending in di-arginine (Fig. S1), are evocative of the Fis1p TA, suggesting a similar, potentially spontaneous mechanism for insertion at mitochondria, although unassisted insertion of the ElaB and YqjD TAs at the mitochondrial surface has yet to be demonstrated. Notably, several conserved members of the current TOM complex are also tail-anchored [30], raising the possibility that at least some of these proteins could be ″hold-overs″ from an early, self-inserting mitochondrial translocon, although we note that these subunits cannot currently self-insert at mitochondria.

Could the DUF883 family of proteins have contributed to an ancestral mitochondrial OM translocon? While YqjD has been reported to recruit ribosomes to the *E. coli* inner membrane during stationary phase [19], a role in line with promotion of co-translational protein import into mitochondria [31,32], the DUF883 family is not readily identified in eukaryotic genomes. One might expect, however, that once a more proficient TOM complex centered around the Tom40 pore evolved, a previous translocon would have been lost, or even selected against if it were to interfere with more rapid protein import through an improved OM translocation machinery. Moreover, an inordinate focus on DUF883 family members when seeking components of the earliest mitochondrial translocon may not be warranted in any case, since the structural characteristics likely required for TA insertion at mitochondria might be easily generated from random open reading frame fragments containing a transmembrane domain. Analogously, random sequences from bacteria are readily able to act as amino-terminal mitochondrial targeting sequences [33-35]. If TAs are easily evolved and might recruit other functional domains to the mitochondrial surface, then identifying orthologs of initial tail-anchored translocon components from existing prokaryotic sequences might be difficult, since an untold number of TAs might be predicted among putative open-reading frames. Supporting the idea that mitochondrial TAs might be generated from sequences not actually functioning in membrane targeting within their native bacterial environment, we demonstrated limited mitochondrial targeting and partial functionality of the computationally predicted TcdA TA in yeast, even though TcdA is unlikely to be membrane-inserted in *E. coli* [36].

If conversion of endosymbiont to mitochondria were the rare and essential event required for generation of eukaryotes, and if insertion of bacteria-derived, tail-anchored proteins at the OM to form an ancestral translocon were necessary for this conversion, then the question of how hospitable an environment the early mitochondria OM might have been for bacteria-derived TAs comes to the fore. Indeed, the membrane into which tail-anchored proteins are inserted can be at least partially determined by their lipid environment [13], and lipids utilized by many characterized archaea are fundamentally different in structure from bacterial and eukaryotic lipids [37]. However, recent evidence indicates that archaeal clades potentially related to the last eukaryotic common ancestor might have been characterized by membranes more similar to those of bacteria than of those membranes more typically found in archaea [38]. This finding raises the possibility that the protoeukaryote’s specific cohort of lipids was crucial to the ability to form complexes of bacteria-derived tail-anchored proteins at the mitochondrial OM that would allow full integration of mitochondria within the ancestral eukaryote.

Finally, we have not examined in detail the trafficking of *E. coli* TAs that appeared to localize to the endomembrane system during our initial survey. However, the diverse organellar locations to which these TAs were localized supports previous data indicating that eukaryotes may derive organelle targeting information from newly acquired prokaryotic proteins or protein fragments, perhaps even from amino acid sequences previously unselected for targeting proficiency [33-35,39,40]. Lateral gene transfer promotes the evolution of novel functions in prokaryotes [41] and was certainly present in the form of endosymbiotic gene transfer during early eukaryogenesis. Indeed, proficiency in making use of cryptic or explicit targeting information in order to direct newly acquired, nucleus-encoded proteins to the distinct subcellular locations where they might be best utilized might have provided a significant selective advantage to the early eukaryote. Such a scenario may be particularly relevant if some amount of cellular compartmentalization already existed in a pre-eukaryotic host cell before conversion of pre-mitochondrial endosymbiont to organelle [42,43].

## CONCLUSIONS

We have demonstrated that TAs from bacteria can localize to and insert within the mitochondrial OM. Our results make plausible the suggestion that tail-anchored proteins acquired by bacteria could have formed an initial translocon at the mitochondrial outer membrane, and our findings indicate that membrane-bound proteins acquired by horizontal gene transfer could have easily found their way to diverse locations within eukaryotic cells at which they might provide a selective advantage. Further efforts will be necessary to determine whether self-inserting proteins or peptides may have generated the initial mitochondrial translocon.

## METHODS

### Yeast strains, plasmids, and culture conditions

Culture conditions are as described in [21], and all experiments have been carried out at 30°C. Strains, plasmids, and oligonucleotides used in this study are found in Supplementary Information 1.

### Selection of *E. coli* tail anchors subject to investigation

FASTA sequences from the *E. coli* proteome were retrieved from UniProt [44] and subjected to analysis using the TMHMM 2.0 server [45]. Polypeptides with a single predicted TM domain (denoted by purple line), harboring 15 or less amino acids carboxyl-terminal to the TM domain, and containing more than 30 amino acids amino-terminal to the TM domain were selected for further analysis.

### Microscopy

Microscopy was performed on logarithmic phase cultures as in [21], with exposure times determined automatically. mCherry fusions are driven by the *ADH1* promoter and universally contain Fis1p amino acids 119-128 (not necessary or sufficient for mitochondrial targeting) linking mCherry to each TA, and genetic assessment of Fis1p variant functionality was performed as described in [21]. The brightness of all images of mCherry expression was adjusted in Adobe Photoshop CS5 (Adobe, San Jose, California) to an equivalent extent. Scoring of mitochondrial morphology was performed blind to genotype.

### Proliferation-based assessment of Fis1p insertion and functionality

Genetic tests of Fis1p insertion and functionality were performed as in [21].

## LIST OF ABBREVIATIONS

TA: tail anchor
mtDNA: mitochondrial DNA
OM: outer membrane
TM: transmembrane
sfGFP: superfolder GFP

## DECLARATIONS

### Ethics approval and consent to participate

None required.

### Consent for publication

None required.

### Availability of data and material

All data generated or analysed during this study are included in this published article and its supplementary information files.

### Authors contributions

C.D.D. designed the study, wrote the manuscript, and performed experiments. G.L.B., A.K., and A.B.S. performed experiments, generated reagents, and provided manuscript critiques. All authors read and approved the final manuscript.

## Acknowledgements

We thank Thomas Richards and Jeremy Wideman for comments on this manuscript.

## SUPPLEMENTAL FIGURE LEGENDS

**Supplemental Figure 1.**
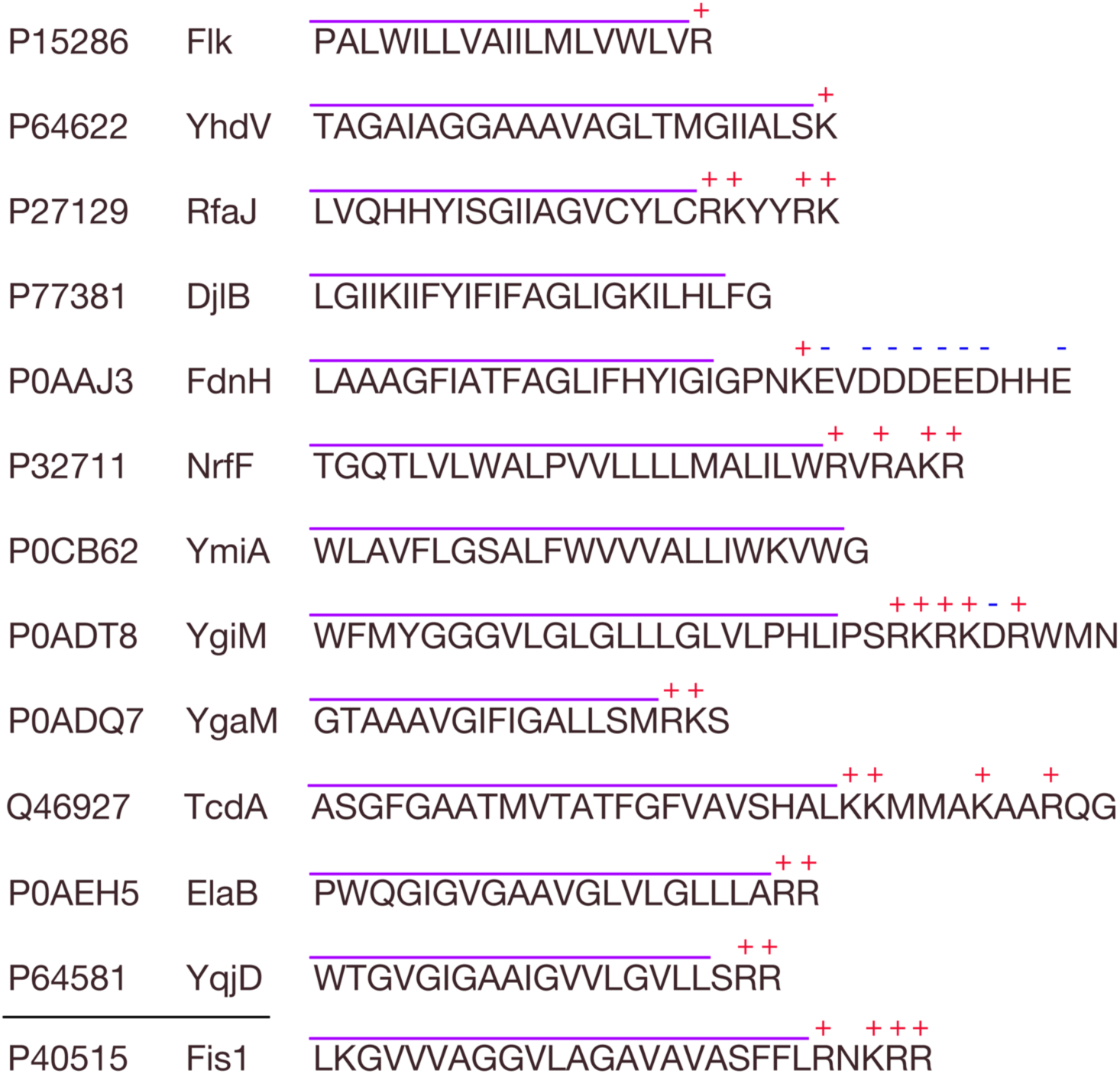
A list of predicted TAs examined in this study. The UniProt accession number and names of selected proteins are provided, along with the sequences of the predicted TAs. Charged amino acids are also denoted. For purposes of sequence comparison, the relevant portion of the *S. cerevisiae* Fis1p TA is also shown.

**Supplemental Figure 2.**
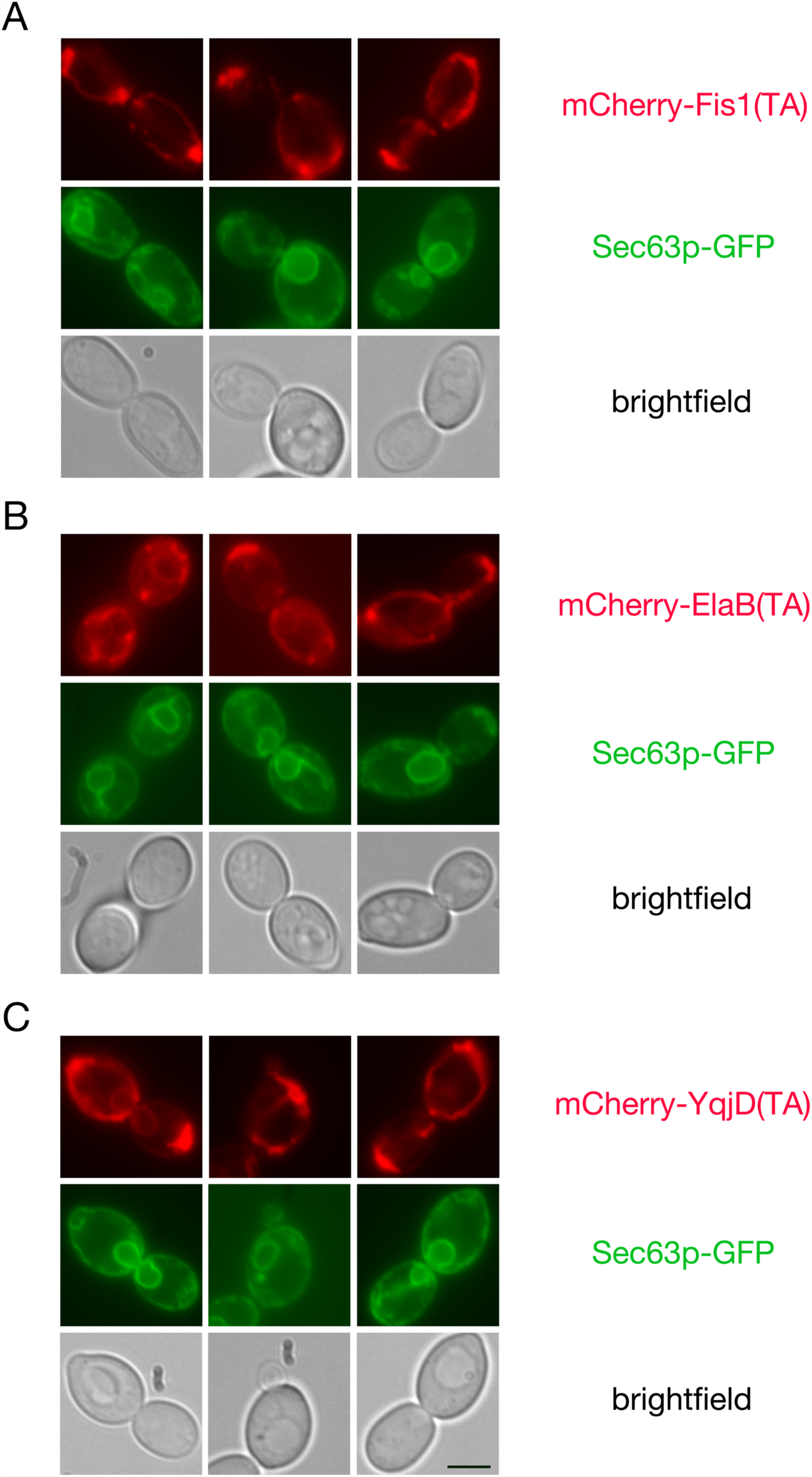
The predicted ElaB and YqjD TAs can also be visualized at the endoplasmic reticulum. Cells were analyzed as in Figure 1, except BY4741 was transformed with plasmid pJK59, expressing Sec63-GFP, before mating.

**Supplemental Figure 3.**
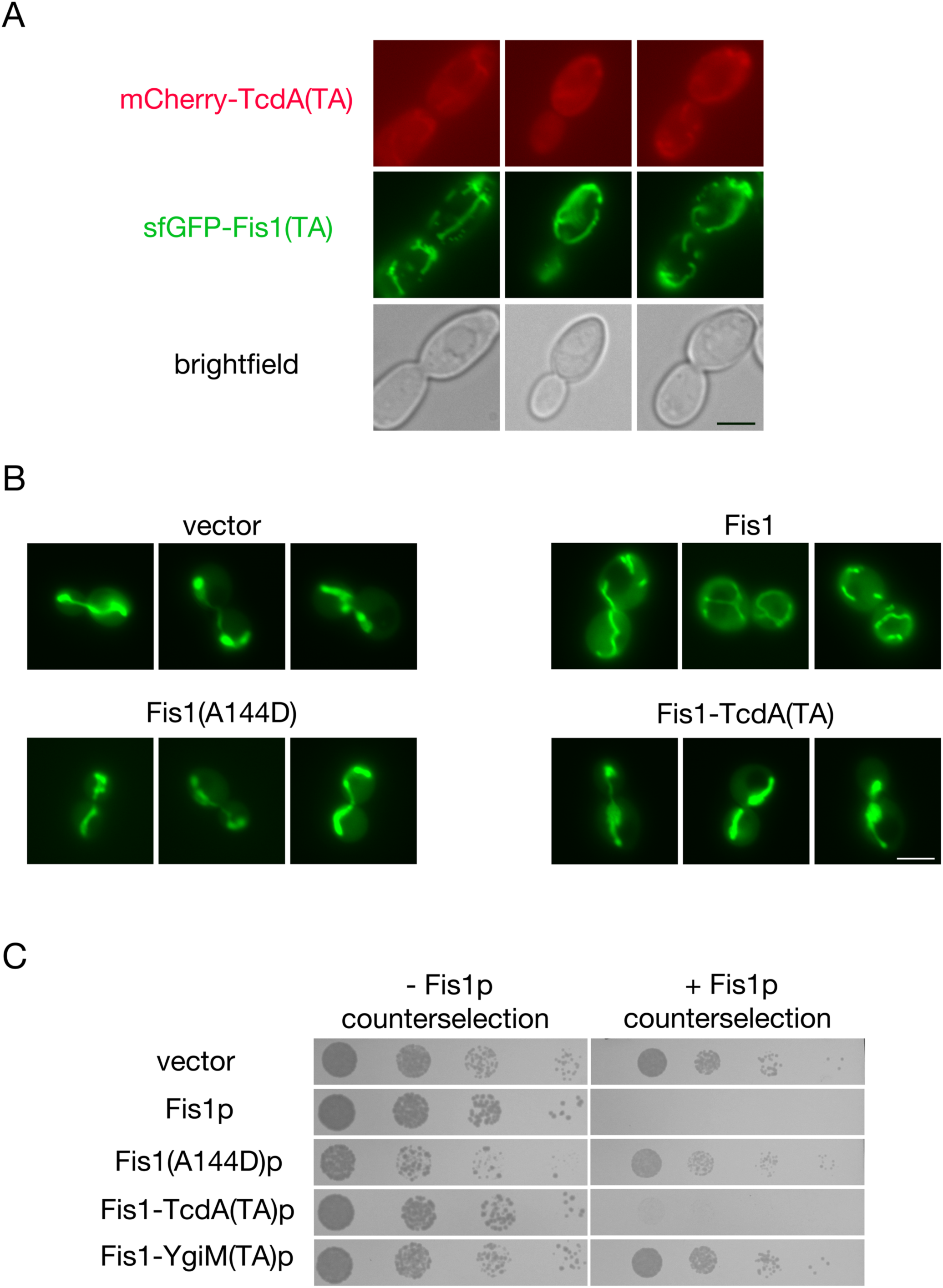
The predicted TcdA TA allows minimal localization to, and function at, the mitochondrial outer membrane. (A) The predicted TcdA TA can be visualized at mitochondria. Strain BY4741, harboring plasmid b294 (sfGFP-Fis1p), was mated to strain BY4742 carrying mCherry-TcdA(TA)-expressing plasmid b281 and the resulting diploids were imaged by fluorscence microscopy. Scale bar, 5 μm. (B) Fis1p with its own TA replaced by the predicted TcdA TA cannot promote normal mitochondrial morphology. *fis1Δ* strain CDD741, expressing mitochondria-targeted GFP from plasmid pHS12, was transformed with empty vector pRS313 or plasmids expressing wild-type Fis1p (b239), Fis1(A144D)p (b244), or Fis1-TcdA(TA)p (b319) and mitochondrial morphology was assessed by fluorescence microscopy. Scale bar, 5μm. (C) Fis1-TcdA(TA)p can allow mitochondrial division. Strain CDD688 was transformed with the plasmids used in (B) or a plasmid expressing Fis1-YgiM(TA)p (b316) and examined as in Fig. 2C, except that culture on medium counter-selective for Fis1p activity was carried out for 5 d.

**Supplemental Figure 4.**
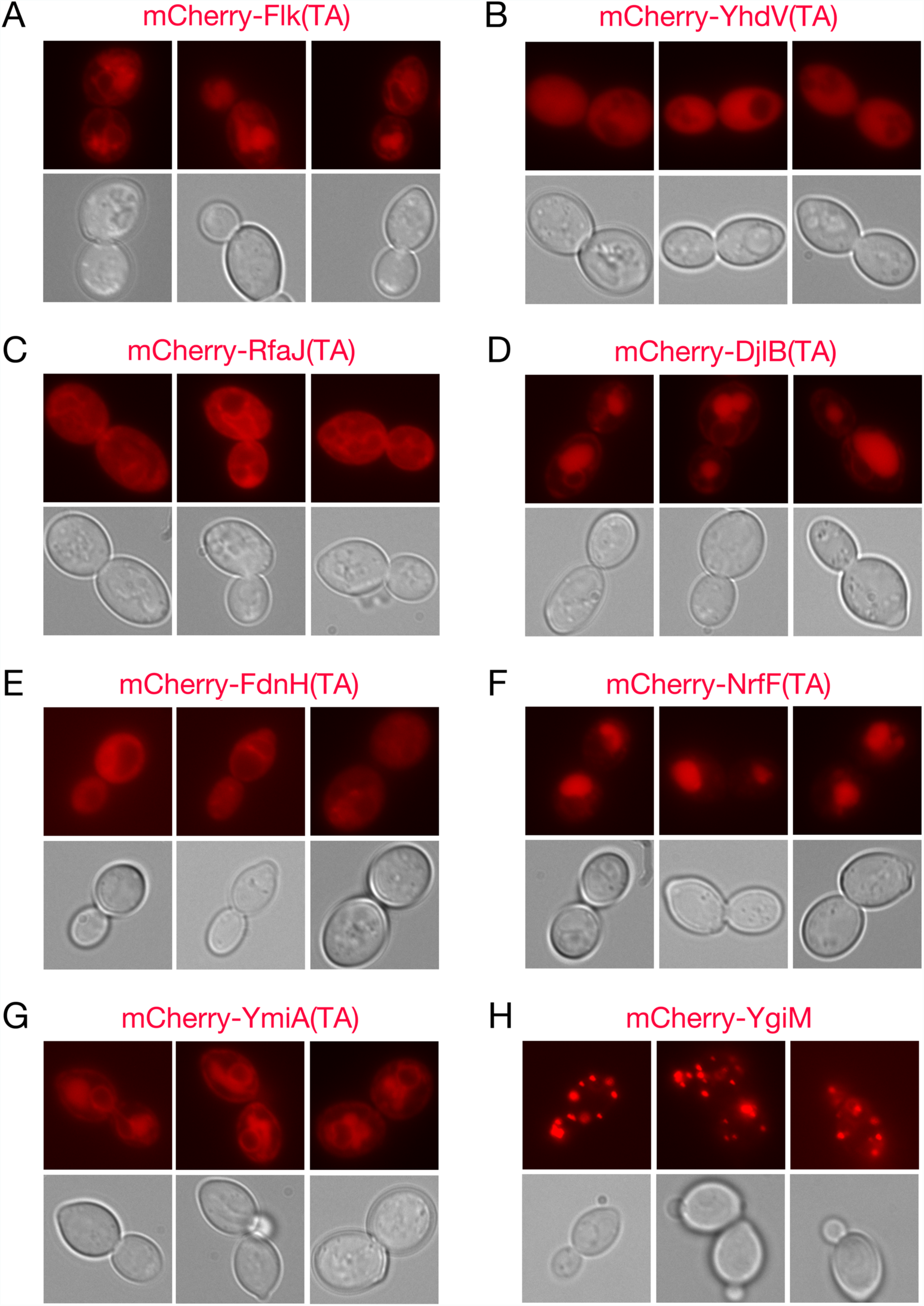
Not all predicted *E. coli* TAs are localized to mitochondria in *S. cerevisiae*. Strain CDD961 was transformed with plasmids expressing (A) mCherry-Flk(TA) (b273), (B) mCherry-YhdV(TA) (b277), (C) mCherry-RfaJ(RA) (b278), (D) mCherry-DjlB(TA) (b280), (E) mCherry-FdnH(TA) (b331), (F) mCherry-NrfF(TA) (b332), (G) mCherry-YmiA(TA) (b333) and examined by fluorescence microscopy. (H) Strain BY4741, carrying plasmid b311 expressing sfGFP fused to the enhanced PTS1 sequence [46], was mated to strain BY4742, containing the mCherry-YgiM(TA)-expressing plasmid b274, and the resulting diploids were imaged.

**Supplementary Information 1. Strains, plasmids, and oligonucleotides used during this study.**

